# Judgments of other bias in Cochrane systematic reviews of interventions are highly inconsistent and thus hindering use and comparability of evidence

**DOI:** 10.1101/366591

**Authors:** Andrija Babic, Andela Pijuk, Lucie Brázdilová, Yuliyana Georgieva, Marco António Raposo Pereira, Tina Poklepovic, Livia Puljak

## Abstract

**Background:** Clinical decisions are made based on Cochrane systematic reviews (CSRs), but implementation of results of evidence syntheses such as CSRs is problematic if the evidence is not prepared consistently. All systematic reviews should assess risk of bias (RoB) in included studies, and in CSRs this is done by using Cochrane RoB tool. However, the tool is not necessarily applied according to the instructions. In this study we aimed to analyze types and judgments of ‘other bias’ in the RoB tool in CSRs of interventions.

**Methods:** We analyzed CSRs that included randomized controlled trials (RCTs) and extracted data regarding ‘other bias’ from the RoB table and accompanying support for the judgment. We categorized different types of other bias.

**Results:** We analyzed 768 CSRs that included 11369 RCTs. There were 602 (78%) CSRs that had ‘other bias’ domain in the RoB tool, and they included a total of 7811 RCTs. In the RoB table of 337 CSRs for at least one of the included trials it was indicated that no other bias was found and supporting explanations were inconsistently judged as low, unclear or high RoB. In the 524 CSRs that described various sources of other bias there were 5762 individual types of explanations which we categorized into 31 groups. The judgments of the same supporting explanations were highly inconsistent. Furthermore, we found numerous other inconsistencies in reporting of sources of other bias in CSRs.

**Conclusion:** Cochrane authors mention a wide range of sources of other bias in the RoB tool and they inconsistently judge the same supporting explanations. Inconsistency in appraising risk of other bias hinders reliability and comparability of Cochrane systematic reviews. Furthermore, discrepant and erroneous judgments of bias in evidence synthesis will inevitably hinder implementation of evidence in routine clinical practice and reduce confidence of practitioners in otherwise trustworthy sources of information.

## Introduction

Assessment of the risk of bias (RoB) in included studies is an integral part of preparing Cochrane systematic reviews (CSRs). Bias is any systematic error that can negatively affect estimated effects of interventions and lead authors to wrong conclusions about efficacy and safety of analyzed interventions [1].

CSRs use Cochrane’s RoB tool, whose aim is to enable better appraisal of evidence and ultimately lead to better healthcare [2]. Cochrane’s standard RoB tool has seven domains, of which first six refer to specific potential biases while the seventh domain is called ‘other bias’, which is used for bias occurring due to any additional problems that were not covered elsewhere in the first six domains [3].

The Cochrane Handbook provides some examples of other potential threats to validity, such as design-specific risk of bias in non-randomized trials, baseline imbalance between groups of participants, blocked randomization in trials that are not blinded, differential diagnostic activity, study changes due to interim results, deviations from the study protocol, giving intervention before randomization, inappropriate administration of an intervention or having co-intervention(s), contamination due to drug pooling among participants, insufficient delivery of intervention, inappropriate inclusion criteria, using instruments that are not sensitive for specific outcomes, selective reporting of subgroups and fraud[3].

This list of potential other sources of bias mentioned in the Cochrane Handbook is limited, and it would therefore be useful to explore potential additional sources of ‘other bias’. By consulting a more comprehensive list of potential other biases, systematic review might recognize certain problems in included studies that might not otherwise consider a potential source of bias.

The aim of this study was to analyze the scoring and support for judgment of the category ‘other bias’ in a large number of interventional CSRs of randomized controlled trials (RCTs) published in the Cochrane Database of Systematic Reviews (CDSR).

## Methods

We conducted a retrospective analysis of published CSRs.

### Inclusion and exclusion criteria

We retrieved CSRs of RCTs about interventions published from July 2015to June 2016 (N = 955) by using Advanced search in The Cochrane Library. Diagnostic CSRs, empty CSRs, overviews of systematic reviews and CSRs withdrawn in this period were excluded. CSRs that included both RCTs and non-randomized trials were included, but only RoB of RCTs was analyzed.

### Screening

One author assessed all titles/abstracts to establish eligibility of CSRs for inclusion. Another author verified the assessments of the first author.

### Data extraction and categorization

Data extraction table was developed and piloted using five CSRs. One author extracted the data and another author verified 10% of extractions. Of the 77 verified CSRs we found 3 CSRs which were partially extracted (3.9%), which we consider to be a negligible percentage of discrepancy. We extracted judgments and supporting explanations for judgments from the other bias section of RoB table in CSRs. We also extracted judgments and support for judgments from additional non-standard domains beyond the seven standard RoB domains in RoB table if Cochrane authors used them. For CSRs that did not use the ‘other bias’ domain in the RoB table or any other additional non-standard domains, we analyzed text of results to see whether Cochrane authors mentioned any potential sources of other bias in the text of the review only. Each supporting explanations for judgments of risk of bias in the analyzed trials was categorized by two authors (AB and LP), via consensus.

### Outcomes

We analyzed number, type, judgments and inconsistencies for various comments about other risk of bias. We also analyzed characteristics of CSRs where there was no ‘other bias’ domain for any of the included RCTs, in terms of number and type of additional non-standard RoB domains that were used instead of ‘other bias’.

### Statistics

We performed descriptive statistics using Microsoft Excel (Microsoft Inc., Redmond, WA, USA). We presented data as frequencies and percentages. In the primary analysis we presented CSRs that had the ‘other bias’ domain in the RoB table. In the secondary analysis we presented CSRs that did not have the ‘other bias’ domain, or had different non-standard variations of risk of bias assessment.

## Results

### 1. Primary analysis

We analyzed 768 CSRs that included 11369 RCTs. Among those 768 CSRs, we included in the primary analysis 602 CSRs that had ‘other bias’ domain in the RoB tables. Those 602 CSRs included a total of 7811 RCTs. We analyzed 166 CSRs in the secondary analysis because they either did not have ‘other bias’ domain in RoB tables (N=149), or those CSRs had both ‘other bias’ domain and additional non-standard domains in the RoB tables (N=17).

Out of 602 CSRs in the primary analysis, there were 524 (87%) CSRs that described various sources of bias in the ‘other bias’ domain, while in 78 (13%) CSRs not a single source of other bias was reported. Furthermore, among 602 CSRs from the primary analysis, there were 337 (56%) CSRs in which at least one included trial indicated that no other bias was found. Terminology for comments about non-existent other bias varied, even within individual CSRs. In 268 (80%) CSRs only one version of the comment that no other bias was found was used, while in 69 (20%) CSRs Cochrane authors used different expressions in comments to indicate that no other sources of bias were found.

In 40 (12%) out of 337 CSRs that indicated that no other bias was found, we observed discrepancies in judgment for this domain. Namely, Cochrane authors in these 40 CSRs sometimes indicated that lack of other bias was associated with low RoB, and sometimes they marked it as unclear or high RoB. In 59 (18%) of these 337 CSRs at least one support for judgment that indicated that no other bias was identified Cochrane authors judged as not being low risk of bias (either high or unclear); in 278 CSRs this was judged as low RoB.

In 19 CSRs all comments that referred to no other bias being identified were judged as unclear. In one CSR having no other bias was judged as both low and high. In one CSR the same comment was judged in different RCTs as either low or high. In one CSR the same comment was judged in different RCTs as either low, or unclear or high.

Of the 7811 trials that were included in the 602 CSRs from the main analysis, in 3703 (47%) trials domain for other bias indicated in the support for judgment that other bias was not identified. Of those 3703 trials, there were 288 (7.8%) that were judged as unclear RoB, 4 (0.1%) that were judged as high RoB, while the others (N=3411, 92.1%) were judged as low RoB.

#### Sources of other bias

In the 524 analyzed CSRs that described various sources of other bias, there were 5762 different supporting explanations for judgments of other bias that we categorized into 31 categories. In 535 trials it was indicated only that it was not possible to assess other bias. For 24 (4%) of those 535 trials it was not indicated why this was not possible, while the most common reasons for not being able to assess other bias were that there were ‘insufficient information’ (N=392, 73%), the trial was published as a conference abstract only (N=78, 15%) and that the trial was published in foreign language so there were issues with translation (N=11, 2%). Cochrane authors were not consistent in judging this type of supporting explanation; for 11 (2%) trials it was judged as high RoB, for 520 (94%) as unclear RoB and for 4 (0.7%) as low RoB.

There were 236 trials for which Cochrane authors simply wrote that issues related to other bias were not described or unclear. This type of supporting explanation was also inconsistently judged by the Cochrane authors; 7 (3%) judged it as low RoB and 229 (97%) as unclear RoB.

The remaining 4991 explanations for judgments of other bias were divided into 29 categories that are shown in Table 1. The most frequently used categories of explanations for other bias were related to baseline characteristics of participants, funding of a trial, reporting, sample size and conflict of interest (Table 2). Cochrane authors used the domain for other bias to indicate positive, negative and unclear aspects of a trial. For example, three most common types of explanations in the category related to baseline characteristic of participants indicated that either baseline characteristics were similar, or that there was imbalance in baseline characteristics, or that there was insufficient information about it. Among 4991 explanations, we were unable to categorize 85 of them because they were uninformative, including explanations such as ‘Adequate’ or ‘N/A’ or ‘Other risk of bias was possible’. Finally, there were 112 explanations that were used only once or twice in RoB tables we analyzed so we categorized that group as ‘Other explanations’.

**Table 1.** Different categories of other bias (based on 4991 explanations) in Cochrane systematic reviews.

| Category | N (%) |
| --- | --- |
| Baseline characteristics of participants | 1067 (21.4) |
| Funding | 774 (15.6) |
| Sample size | 405 (8.1) |
| Reporting | 381 (7.6) |
| Conflict of interest | 288 (5.8) |
| Inclusion and exclusion criteria | 197 (3.9) |
| Confounding | 196 (3.9) |
| Analyses | 191 (3.8) |
| Outcome domains and outcome measures | 135 (2.7) |
| Co-interventions | 134 (2.7) |
| Deviations from the protocol | 123 (2.5) |
| Randomisation | 111 (2.2) |
| Terminated early | 108 (2.2) |
| Issues related to cross-over trials | 98 (2) |
| Intention-to-treat analysis (ITT) | 95 (1.9) |
| Study design | 76 (1.6) |
| Compliance | 72 (1.4) |
| Attrition | 71 (1.4) |
| Contamination | 65 (1.3) |
| Follow-up and study duration | 46 (0.9) |
| Blinding | 25 (0.5) |
| Clustering | 17 (0.3) |
| Selection bias | 17 (0.3) |
| Protocol registration | 16 (0.3) |
| Study quality | 9 (0.2) |
| Publication bias | 7 (0.1) |
| Adequacy of comparators | 5 (0.1) |
| Inexplicable | 85 (1.7) |
| Other | 177 (3.6) |

#### Partial studies included in the primary analysis

We found 34 CSRs with specific partial data regarding other bias. We divided them into four distinct groups: first group with 28 CSRs that had judgments for ‘other bias’, but not all had accompanying comments, second group with 4 CSRs where only one included RCT did not have the ‘other bias’ domain, third group with one CSR with included RCT without ‘other bias’ domain and included RCT with only judgment without comment, and fourth group with one CSR where RoB table was completely missing for 6 included RCTs. Some CSRs had additional non-standard RoB domains, separately or in addition to the ‘other bias’ domain. Categories of additional non-standard RoB domains in CSRs are shown in Table 3.

**Table 2.** Judgments for the 20 most common explanations of other bias.

| **Explanation** | **Total** | **High, N (%); n\*** | **Unclear, N (%); n\*** | **Low, N (%); n\*** |
| --- | --- | --- | --- | --- |
| Not possible to assess other bias | 504 | 7 (1.4);7 | 494 (98);117 | 3 (0.6);3 |
| Baseline characteristics similar between the groups | 314 | 0 (0);0 | 24 (8);13 | 290 (92);61 |
| Not described/unclear | 233 | 0 (0);0 | 226 (97);54 | 7 (3);4 |
| Baseline imbalance between groups of participants | 167 | 91 (54);56 | 62 (37);41 | 14 (9);12 |
| Funding: industry | 162 | 83 (51);28 | 77 (48);25 | 2 (1);2 |
| Potential confounding factors | 120 | 63 (53);38 | 47 (39);34 | 10 (8);9 |
| Not enough information on baseline characteristics of participants | 88 | 8 (9);6 | 78 (89);39 | 2 (2);2 |
| Funding: non-profit | 86 | 0 (0);0 | 4 (5);4 | 82 (95);33 |
| Funding: not reported | 72 | 0 (0);0 | 68 (94);15 | 4 (6);4 |
| Important parameters not reported | 61 | 19 (31);14 | 41 (68);28 | 1 (1);1 |
| Sample size: calculation of sample size not provided | 42 | 24 (57);6 | 17 (41);7 | 1 (2);1 |
| Potential randomisation problem | 40 | 9 (23);9 | 28 (70);13 | 3 (7);3 |
| Potential problem with inclusion criteria | 40 | 16 (40);15 | 22 (55);12 | 2 (5);2 |
| Deviations from the study protocol | 37 | 16 (43)<br>13 | 18 (49)<br>15 | 3 (8)<br>3 |
| No relevant subgroup analysis | 36 | 10 (28);1 | 26 (72);1 | 0 (0);0 |
| Funding: intervention supplied by industry | 32 | 14 (44);7 | 12 (38);10 | 6 (18);3 |
| Adequate | 28 | 0 (0);0 | 0 (0);0 | 28 (100);1 |
| No information on the validity of the outcome measure | 27 | 3 (11);3 | 23 (85);5 | 1 (4);1 |
| Sample size: performed calculation | 24 | 1 (4);1 | 3 (12);3 | 20 (84);9 |
| Sample size: small | 23 | 8 (35);5 | 15 (65);5 | 0 (0);0 |

\*n= Number of SRs that included at least one RCT with this characteristic

**Table 3.** Categories of additional non-standard RoB domains in Cochrane systematic reviews.

| **Additional category** | **N of CSRs** |
| --- | --- |
| Group similarity at baseline (selection bias) | 11 |
| Baseline data | 5 |
| Baseline outcome measures (similar) | 3 |
| Groups balanced at baseline/ balance in baseline characteristics | 2 |
| Baseline characteristics of participants | 1 |
| Baseline comparability of treatment and control groups | 1 |
| Baseline measures | 1 |
| Similarity of baseline characteristics* | 1 |
| Treatment/control groups comparative at entry | 1 |
| Major imbalance in important baseline confounders | 1 |
| Comparability of groups on different prognostic characteristics* | 1 |
| Size | 8 |
| Size of the study | 5 |
| Small sample size bias | 4 |
| Sample size* | 2 |
| Sufficient sample size* | 1 |
| Power calculation* | 1 |
| Timing of outcome assessment (similar)* | 10 |
| Adequate follow-up | 2 |
| Study duration | 2 |
| Early stopping | 1 |
| Groups received comparable treatment | 2 |
| Care program identical/ identical care | 2 |
| Treatment fidelity* | 1 |
| Free of systematic differences in care?* | 1 |
| Consistency in intervention delivery | 1 |
| Equality of treatment | 1 |
| Protocol deviation balanced | 1 |
| Groups received same intervention | 1 |
| Compliance/adherence assessed (acceptable) | 7 |
| Compliance with recommendation reliable? | 1 |
| Compliance acceptable* | 1 |
| Source of funding/ sponsorship | 4 |
| For profit funding* | 1 |
| Funding* | 1 |
| Vested interest bias | 1 |
| Conflict of interest | 1 |
| Co-intervention avoided or similar* | 5 |
| Co-interventions | 2 |
| Groups received same co- interventions | 1 |
| Intention to treat | 5 |
| Incorrect analysis | 1 |
| Results based on data dredging? | 1 |
| Analyses adjust for different lengths of follow-up workers? | 1 |
| Appropriate statistical tests use? | 1 |
| Adequate adjustment for confounding in the analyses? | 1 |
| Contamination/ protection against contamination | 3 |
| Validity of outcome measures | 1 |
| Reliability of outcome measures | 1 |
| Outcome measures used valid and reliable? | 1 |
| Free from performance bias | 1 |
| Performance bias as «differential expertise» bias | 1 |
| Performance bias as comparability in the experience of care providers | 1 |
| Adequate patient description | 1 |
| Recruitment of participants from the same population? | 1 |
| Recruitment of participants over the same study period? | 1 |
| Washout/ carry-over effect in cross-over study designs | 2 |
| Overall assessment of bias risk | 1 |
| Summary of risk of bias for Consumption outcome | 1 |
| Researcher allegiance* | 1 |
| Therapist allegiance* | 1 |
| CHBG (Cochrane hepato-biliary group) combined assessment (mortality)* | 1 |
| CHBG combined assessment (hepatic encephalopathy)* | 1 |
| Comparability with individually randomized trials | 1 |
| Detection bias (biochemical validation of smoking outcomes) | 1 |
| Ethical approval | 1 |
| Explicit inclusion/exclusion criteria | 1 |
| Free of dietary differences other than fat?* | 1 |
| Loss of clusters | 1 |
| Methods for selecting cases to adjudicate | 1 |
| Outcome description | 1 |
| Publication format | 1 |
| Recruitment bias | 1 |

\*domains found in 9 Cochrane reviews that had both ‘other bias’ domain and additional non- standard domain(s) for other bias in RoB tables

#### Cochrane authors’ judgments of different explanations for ‘other bias’

There were 3033 trials for which only one category of explanation was written by Cochrane authors. When the explanation had only one category of comment we could be certain that the judgment referred only to that specific comment so we analyzed those in detail to see how the Cochrane authors judge different explanatory comments. There were 259 types of different explanations among those 3033 trials. We analyzed in more detail those judgments for 20 most common explanations of other bias and found very high inconsistency in how Cochrane authors judge the same explanations (Table 2).

### 2. Secondary analysis

#### Reviews without ‘other bias’ domain in the RoB table

Among 149 CSRs that did not have ‘other bias’ domain in the RoB table, there were 102 CSRs that did not have any other replacement domain for ‘other bias’. These 102 CSRs used varied number of standard RoB domains. In those 102 CSRs, number of standard RoB domains that were used varied, with one standard RoB domain in 4 CSRs, three RoB domains in 7 CSRs, four RoB domains in 15 CSRs, five domains in 51 CSRs and 6 domains in 25 CSRs.

For this group of CSRs, that did not have the ‘other bias’ domain in the RoB table, we analyzed texts of results to see whether they mentioned any other sources of bias, beyond the standard six domains, in the section ‘Risk of bias in included studies’. We found that 68/102 (67%) did not mention any sources of other bias in the results of review. However, the remaining 34 (33%) did have comments about the other bias. Three of those 34 stated that they had not found any other risk of bias, while 31 CSRs out of those 34 reported in the text of results that the included studies had had from 1 to 6 different categories of other bias.

#### Reviews with both ‘other bias’ domain and additional non-standard domain(s) for other bias in RoB tables

Nine CSRs had both ‘other bias’ domain and additional non-standard domain(s) for other bias in RoB tables. Those CSRs used from 1 to 4 additional non-standard domains; 18 in total. Those additional non-standard RoB domains are listed in Table 3 and marked with asterisk.

#### Reviews without ‘other bias’ domain but with additional non-standard domain(s)

There were 57 CSRs that did not have the ‘other bias’ domain, but they did have additional non-standard RoB domains apart from the standard domains in the Cochrane RoB table. Most of the CSRs had only one additional non-standard domain (N=24), while others had 2-8 additional domains per each RCT. Table 3 shows non-standard domains that were used in those CSRs without ‘other bias’ domain.

#### Reviews that consistently did not use support for judgment or they used non-standard judgments

We found 9 CSRs that consistently did not use supporting explanations for judgment or they used non-standard judgments. In 5 CSRs authors used judgments low, high or unclear RoB, but without comments as support for judgment. In one CSR all trials were marked with unclear risk of other bias without any comment as support for judgment. In four CSRs all trials were marked with low risk of other bias without any comment as support for judgment. We also found 4 CSRs that did not have judgments low-high-unclear, but different kinds of judgments. One CSR had judgments yes/no without supporting comments; two CSRs had judgments yes, no or unclear, with supporting comments and there was one CSR with judgments A-adequate and B-unclear.

## Discussion

In this study we analyzed 768 Cochrane systematic reviews, with 11369 included trials. We found that Cochrane authors used numerous different categories of sources of other bias and that they were not judging them consistently. We categorized different types of supporting explanations into 31 categories, and we found numerous other inconsistencies in reporting of sources of other bias in CSRs. Findings of this study are disconcerting because consistency in secondary research is very important to ensure comparability of studies.

Insufficient and unclear reporting of the ‘other bias’ domain was very common in the CSRs we analyzed. Among the most common support for judgment were comments that we categorized as ‘not described/unclear’, which is puzzling because ‘other bias’ domain is not specific like the other six domains of the RoB tool, and it is therefore difficult to fathom what it means that other bias was not described or that it was unclear. If the authors did not find sources of other bias, or if they thought that they could not assess other bias because of brevity of report or language issues, they should have stated that. Likewise, for some trials the only supporting explanation was that other bias was ‘Adequate’. Without any further explanations, readers cannot know what exactly the Cochrane authors found to be adequate in terms of other potential sources of bias. Many systematic reviews had a high number of included studies, and therefore some comments were repeated multiple times in the same systematic review.

The most commonly used specific category of other bias referred to baseline characteristics of participants. In RCTs randomization should ensure allocation of participants into groups that differ only in intervention they received. Randomization should ensure that characteristics of participants that may influence the outcome will be distributed equally across trial arms so that any difference in outcomes can be assumed to be a consequence of intervention [4].Baseline imbalances between the groups may indicate that there was something wrong with the randomization process, or that they might be due to chance [5]. Severe baseline imbalances can occur because of deliberate actions of trialists if they aim to intentionally subvert the randomization process [6]or due to unintentional errors.

Chance imbalances should not be considered a source of bias, but it may be difficult to distinguish whether baseline imbalances are caused by chance or intentional actions. If there are multiple studies included in a meta-analysis, it could be expected that chance imbalances will act in opposite directions. But the problem may occur if there is a pattern of imbalances across several trials that may favor one intervention over another, suggesting imbalance due to bias and not due to chance [7]. Cochrane is now developing a second generation of the RoB tool, titled RoB 2.0, and one of the signaling questions in the RoB domain about randomization process asks “Were there baseline imbalances that suggest a problem with the randomization process”[7]. The fact that so many Cochrane authors used comments about baseline imbalance as a domain of other bias, and not in the RoB domain about random sequence generation (selection bias) indicate that many Cochrane authors consider that this aspect should be emphasized separately from the selection bias domain.

The second most commonly used category of supporting explanations was related to funding of a trial, and comments about conflicts of interest were the fifth most common category. This is in direct contrast with the recommendations from the Cochrane Handbook, where it is acknowledged that information about vested interests should be collected and presented when relevant, but not in the RoB table; such information should be reported in the table called ‘Characteristics of included studies’ [8]. RoB table should be used to describe specific methodological aspects that may have been influenced by the vested interest and directly lead to RoB [8]. Therefore, it is obvious that the authors of the Cochrane Handbook assume that the influence of sponsors can be mediated via other domains of RoB tool such as selective reporting of favorable outcomes.

However, Lundh et al. have published a CSR in 2017 about industry sponsorship and research outcomes, in which they included 75 primary studies, which shows that commercial funding leads to more favorable efficacy results and conclusions compared to non-profit funding [9]. They concluded that industry sponsorship introduces bias that cannot be explained by standard domains of Cochrane’s RoB assessment [9]. The debate about whether funding presents source of bias or not is ongoing in the Cochrane, with some considering that commercial funding is a clear risk of bias, while others argue against such standpoint[10, 11]. This debate apparently reflects the current situation in which many Cochrane authors continue to use funding and conflict of interest as a source of other bias despite the official warning against such use of information about sponsorship from the Cochrane Handbook, as we have demonstrated in this study.

The third most frequent category of supporting explanations for other bias was related to poor reporting, where Cochrane authors indicated that relevant information were missing or were inadequately reported. Poor reporting hinders transparency, as it allows authors to avoid attention to weak aspects of their studies. For this reason reporting guidelines should be used [12].

Comments about sample size were the fourth most common category either in a sense that the trial did or did not report sample size calculation, or that sample size was “small” without any further explanation of what the Cochrane authors considered to be a small sample. There were 21 trials for which Cochrane authors wrote that there were fewer than 50 participants in each arm. It is unclear where this cut-off is coming from, as there is no such guidance in the Cochrane Handbook in the chapter about risk of bias. On the contrary, chapter 8.15.2. of the Cochrane Handbook specifically warns that “sample size or use of a sample size (or power) calculation” are examples of quality indicators that „should not be assessed within this domain”[8].

The Cochrane Handbook also warns that authors should avoid double-counting, by not including potential sources of bias in the ‘other bias’ domain if they can be more appropriately covered by other domains in the tool [8]. As can be seen by our study, Cochrane authors sometimes do double-counting because there were categories of comments supporting judgments that could have been addressed in the first six domains.

As we have shown, most of Cochrane authors decided to use the other bias domain to describe potential additional biases that were not covered in the first six domains of the RoB tool. In the proposed RoB tool 2.0 there is no ‘other bias’ domain [7]. The proposed RoB tool is much more complex, compared to the current version of the RoB tool, and many items that were specifically emphasized by Cochrane authors in the other bias domain, as shown in our study, are addressed in the RoB 2.0 tool. However, there are still potential biases from other sources that the RoB 2.0 may neglect by omitting the RoB domain, such as biases specific to certain topics, and those that were not recognized by the RoB 2.0 tool in advance.

We have already conducted a similar analysis of Cochrane RoB domain related to attrition bias, and we found that judgments and supports for judgments in that domain were extremely inconsistent in CSRs (unpublished data). This analysis related to sources of other bias in CSRs contributes to the perception that Cochrane RoB tool is inconsistently used among Cochrane authors. The authors do not necessarily follow guidance from the Cochrane Handbook. In the support for judgment they mention issues that the Cochrane Handbook explicitly warns against. Various comments that serve as supports for judgments were inconsistently judged across CSRs and trials included in CSRs. Cochrane authors also use inconsistent terminology to describe the same concepts. Increasing complexity of the RoB tool, as proposed in the RoB tool 2.0 will likely only increase this problem of insufficient consistency in RoB appraisal and worsen this problem of insufficient comparability of judgments of RoB across CSRs.

Furthermore, our study indicated that Cochrane authors extensively use the available option to customize the RoB table. We found that there were as many as 102 (13%) out of 768 analyzed CSRs that did not use the other bias domain in the RoB table at all. CSRs are produced using software Review Manager (RevMan). As soon as an author inserts a new study in the RevMan among included studies, an empty RoB table for the study automatically appears, with seven pre-determined domains. Therefore, Cochrane authors need to intentionally remove or add some domains if they want to customize the RoB table. Among 102 CSRs that did not have other bias domain, 33% of those CSRs had comments about other potential sources of bias in the body of the manuscript. It is unclear why some Cochrane authors use only text for comments about other bias instead of using RoB table for this purpose. Additionally, we observed that in many CSRs without other bias domain there were other customizations of the RoB table, which had from one to six other, standard RoB domains included. Exactly half of those CSRs without other bias domain in the RoB table had less than six standard domains in the RoB table.

Limitation of our study is a limited number of analyzed CSRs that were published in 2015 and 2016. However, considering the number of CSRs analyzed, and the amount of inconsistency we observed, we have no reason to suspect that the results would be significantly different if a more recent cohort of published CSRs would have been used. Additionally, it takes a long time to categorize thousands of different inconsistent supporting explanations. Some unintentional errors in categorizations may have been made.

## Conclusion

Cochrane authors mention a wide range of sources of other bias in the RoB tool and they inconsistently judge the same supporting explanations. Inconsistency in appraising risk of other bias hinders reliability and comparability of Cochrane systematic reviews. Discrepant and erroneous judgments of bias in evidence synthesis will inevitably hinder implementation of evidence in routine clinical practice and reduce confidence of practitioners in otherwise trustworthy sources of information. Potential remedies include more attention to author training, better resources for Cochrane authors, better peer-review and editorial consistency in the production of Cochrane systematic reviews.

## Declarations

### Ethics approval and consent to participate

Not applicable; this was secondary study

### Consent for publication

Not applicable

### Availability of data and material

The datasets used and/or analysed during the current study are available from the corresponding author on reasonable request.

### Competing interests

The authors declare that they have no competing interests.

### Funding

No extramural funding.

### Authors’ contributions

Study design: LP. Data analysis and interpretation: AB, AP, LB, YG, MARP, TPP, LP. Drafting the first version of the manuscript: AB, LP. Revisions of the manuscript: AB, AP, LB, YG, MARP, TPP, LP. All authors read and approved the final manuscript. All authors agree to be accountable for this work.

## Acknowledgements

We are grateful to Ms. Dalibora Behmen for language editing.

